# Compositional forecasting of Chinook Salmon Evolutionarily Significant Units in bycatch for Pacific Hake fisheries

**DOI:** 10.1101/2021.11.29.470462

**Authors:** Paul Moran, Vanessa J. Tuttle, Susan Bishop, Larrie LaVoy

## Abstract

Bycatch impacts on non-target species present significant management problems in diverse fisheries throughout the world. Despite successful efforts to minimize bycatch in US West Coast Pacific Hake fisheries, these impacts remain a concern, particularly for sensitive populations of Chinook Salmon. NOAA Fisheries needed predictive models to estimate proportions of Chinook Salmon Evolutionarily Significant Units (ESUs) expected in bycatch. We used genetic mixture analysis to estimate ESU proportions from at-sea bycatch between 2008 and 2015. Using latitude as a predictor and applying jackknife cross validation, we found Dirichlet regression more accurately estimated abundant ESUs, whereas multinomial logistic regression performed better with rare ESUs. This targeted, ESU-specific approach showed the spatial distribution of sensitive stocks in bycatch and supported NOAA’s obligations to forecast impacts on listed ESUs. The overarching goal of this continuing work is to maximize sustainable harvest while protecting threatened and endangered Chinook Salmon ESUs.

## INTRODUCTION

Pacific Hake^1^ (*Merluccius productus*) are distributed from the Gulf of Alaska to the Gulf of California (Quirollo et al. 2001). This abundant marine resource supports a large-scale trawl fishery off the US West Coast. Trawling in this region began slowly in the 1870s but increased in the 1920s with implementation of diesel engines and other technological advances (Easley 2001). By 1966 harvest reached 137 kMT and now represents an important economic resource to the region and to the nation (Hamel et al. 2015). In 2016, commercial landings of Pacific Hake totaled more than 260.8 kMT valued at over US$42 million (NOAA 2019).

### Incidental Take Statement for the US West Coast groundfish fisheries

Despite much effort and management action to reduce impacts on non-target species, bycatch remains a concern in this commercially important fishery (Somers et al. 2015). There is special concern for bycatch of Chinook Salmon (*Oncorhynchus tshawytscha*) protected under the US Endangered Species Act (ESA). By international treaty (PSC 2020), management of coastal Pacific hake fisheries is shared among Canadian Department of Fisheries and Oceans, NOAA Fisheries, and the Joint Management Committee. In cooperation with the Pacific Fishery Management Council (PFMC), NOAA manages the Pacific Hake fishery on the US West Coast in Federal waters (3 to 200 miles offshore). Federal agencies must consult with NOAA on activities that might jeopardize the continued existence of protected marine species (essentially any action that involves “take”). As part of the 2017 ESA Consultation for the West Coast Groundfish Fishery Management Plan salmon biological opinion (BiOp), the PFMC and NOAA faced a series of questions related to alternative fishery regulation scenarios. For example: What would be the actual number of Chinook Salmon individuals in bycatch from each Evolutionarily Significant Unit (ESU), if the current restriction were rescinded on processing Pacific Hake south of latitude 42? What if there was a resumption of the tribal mothership fishery off the north coast of Washington State? Ultimately, NOAA’s Incidental Take Statement needed to forecast the actual number of Chinook Salmon from each ESU taken in bycatch, given predicted spatial distribution of fishing effort (Matson and Erickson 2018) and under different management scenarios (NMFS—WCR 2017).

### Limited CWT recoveries in bycatch

Tiny coded-wire tags (CWTs) are implanted in the snouts of juvenile hatchery fish, and much has been learned about Chinook Salmon migration patterns and ocean distribution from CWT recoveries (Weitkamp 2010; Riddell et al. 2018; Shelton et al 2019). We know that particular stocks have characteristic patterns of tag recovery, primarily in commercial salmon harvest. Importantly, these patterns of migration were shown to be surprisingly stable across years, despite high interannual variation in ocean conditions and relative abundance (Weitkamp 2010). In spite of broad utility in harvest management and basic research, coded-wire-tag recoveries in bycatch have generally been inadequate to estimate relative abundance and distribution of 17 ESUs (identified in Appendix 1). In parallel with tissue sampling, only 687 CWTs were recovered from among 9,862 Chinook Salmon sampled by NOAA At-Sea Observers over the 8 years of this study (2008 – 2015). Therefore, genetic analysis, where every fish, hatchery and wild, is effectively “tagged,” presented an opportunity to significantly augment what was known from sparse CWT recoveries (NMFS—WCR 2017).

In the current study, we used genetic mixture analysis to characterize stock composition of Chinook Salmon ESUs in the US West Coast, at-sea, Pacific Hake fishery. We also tested accuracy and precision of different predictive regression models used to forecast ESU impacts in a management context; the goal being to provide the most powerful forecasting tool requiring the simplest possible inputs. Beyond specific interest to salmon conservation and Pacific hake harvest, our results are relevant to broader studies of more general statistical challenges of multinomial regression modeling and measures of forecasting accuracy—both common problems in ecology and natural resource management.

## MATERIALS AND METHODS

NOAA’s At-Sea Hake Observer Program (A-SHOP) collected Chinook Salmon tissue samples from the catcher/processor, mothership, and at-sea tribal sectors (NWFSC 2021). Rayed fin-clip samples were folded in Whatman 3MM chromatography paper, dried immediately, and stored in barcoded coin envelopes at ambient temperature. Samples were deposited in the Northwest Fisheries Science Center (NWFSC) Conservation Biology Division’s Genetic Tissue Archive (accession numbers in Table 1). All samples were collected during the normal fishing season beginning 15 May and ending 31 December. Tissue samples selected for genotyping (4,498) were drawn randomly each year between 2008 and 2015. Annual sample sizes were dictated by project resources available in a given year, not total bycatch. Table 1 illustrates the filtering process and identifies the following: total estimated bycatch, samples collected by observers, random sub-sample for genotyping, and filtered for genotyping quality and filtered for ≥0.8 assignment probability. The latter two sample sets provided the primary input data for this study and reflect two distinctly different statistical approaches; 1) modeled genetic stock composition estimates based on all individuals for all ESUs simultaneously, and 2) discrete individual assignment of each fish to population of origin. Our sample was intended to accurately reflect ESU-specific, spatial and temporal bycatch impacts over the course of each year in the fishery. That was the essential focus of NOAA’s BiOp as related to Chinook salmon impacts (NMFS—WCR 2017).

**Table 1.** Chinook Salmon bycatch in the at-sea sectors of the US West Coast Pacific Hake fishery (where >0.8 is the number of individual fish that met that assignment probability threshold, and Mean latitude is derived from all fish genotyped)

| **Year** | **Bycatch<sup>1</sup>** | **Collected** | **Subsampled** | **Genotyped** | **>0.8** | **Mean lat.** | **Accession #<sup>2</sup>** |
| --- | --- | --- | --- | --- | --- | --- | --- |
| **2008** | 875 | 271 | 271 | 258 | 178 | 46.32 | 34801 |
| **2009** | 1,142 | 403 | 403 | 390 | 326 | 47.89 | 34592 |
| **2010** | 1,364 | 680 | 680 | 664 | 548 | 46.89 | 34927 |
| **2011** | 4,360 | 1,837 | 1,835<sup>3</sup> | 1,693 | 1,284 | 44.77 | 90517 |
| **2012** | 4,209 | 2,013 | 288 | 288 | 215 | 44.43 | 90598 |
| **2013** | 3,739 | 1,542 | 288 | 285 | 218 | 43.78 | 90601 |
| **2014** | 6,695 | 2,368 | 384 | 381 | 295 | 43.54 | 90627 |
| **2015** | 1,806 | 748 | 349 | 345 | 296 | 43.91 | 90654 |
| **Total** | 24,190 | 9,862 | 4,498 | 4,304 | 3,360 | 45.21 |  |

<sup>1</sup>Estimated total Chinook bycatch
nwfsc.noaa.gov/research/divisions/fram/observation/xls/SalmonBycatch_Expanded_2002-2015.xlsx
<sup>2</sup>NWFSC Conservation Biology Division Tissue Archive
<sup>3</sup>Essentially all collected samples were selected for genotyping in 2012

### Genotyping, genetic mixture modeling, and individual assignment

DNA was extracted and purified by using Qiagen® DNeasy™ membrane capture. Purified DNA was amplified and genotyped for 13 internationally standardized microsatellite loci (see below). Microsatellite products were sized using an Applied Biosystems Incorporated (ABI) 3100 Genetic Analyzer. Genotypes were inferred from electropherograms by using ABI Genescan and Genotyper software. We used conditional maximum likelihood mixture modeling (CMLMM) to simultaneously estimate stock compositions and make individual assignments to population of origin (Rannala and Mountain 1997; ONCOR, Kalinowski et al. 2007), with bias correction (Anderson et al. 2008). Population-level allocation was then aggregated by ESU reporting group. Genetic mixture analysis using ONCOR was replicated and confirmed with the R package ‘rubias’ (Moran and Anderson 2019). The two sets of genetic results from ONCOR, 1) modeled proportions and 2) individual assignments, were used for the two different classes of statistical analysis, 1) Dirichlet regression (DR; Maier 2014) and 2) multinomial logistic regression (MLR; Hilbe 2009). CMLMM is taken to be the best possible estimate of ESU proportions in a given year. We use the term “observed” in reference to observed genotypes and observed latitudes that are compared to “predicted” ESU proportions from the regression models. We calculated credibility intervals for CMLMM proportions, however, we relied principally on cross validation of predictions and evaluation of accuracy with scale-dependent and -independent metrics (see below).

CMLMM depends on a baseline dataset of known-origin reference samples that is assumed to represent all potentially contributing populations. In this study, we used the most comprehensive Chinook Salmon baseline available, the internationally standardized, microsatellite dataset (i.e., common loci and consistent allele designations, Moran et al. 2006) that was developed by the Genetic Analysis of Pacific Salmonids consortium (GAPS; Moran et al. 2005; Seeb et al. 2007). The GAPS baseline was designed for eastern Pacific fishery mixtures, primarily harvest, but the geographic coverage of potential source populations is complete from Southeast Alaska to Central Valley, California. The version of the baseline we used included more than 20,000 known-origin fish from 163 populations representing all ESUs and major Canadian and Alaskan stocks that contribute to these fisheries (Appendix 1). The GAPS baseline is thoroughly vetted by the salmon genetics community and has been widely used in studies of harvest impacts (Bellinger et al. 2015; Satterthwaite et al. 2015; Moran et al. 2018).

### Predictive regression models and cross validation

Input data for our preliminary exploration included the ESU to which each fish assigned, collection year, ordinal day of the year, latitude (decimal degrees), fork length (cm), and fishing depth (m). Akaike information criterion (AIC) was used to evaluate different predictive models. Focusing on mean latitude of an annual bycatch sample, we compared two multinomial regression methods for predictive forecasting. DR was used to relate observed proportions for an annual sample of individuals relative to the mean latitudes over which those individuals were taken. MLR was also used to estimate an expected proportion for each ESU as a function of latitude but was based on individual fish rather than annual means. Compositional proportions are bounded by zero and one, and not normally distributed or homoscedastic. This class of proportions violates assumptions of linear regression and common parametric analyses such as t-tests and ANOVA, when used as a dependent variable. To evaluate the absolute and relative accuracy and precision of the two regression methods, we conducted sequential, independent, cross-validation analyses holding out each of 8 individual years, one at a time, as test datasets and using all remaining years as training data. We used a similar jackknife approach for both regression methods. In each case, we evaluated the accuracy of the prediction (based on the training data set) against the independent, cross-validation, test set (the “actual” or “observed” ESU proportions derived from observed genotypes). Central Valley Spring ESU only occurred in one test set (annual sample), so that ESU was omitted from the jackknife. To the extent possible, the jackknife is a sequential test of forecasting accuracy, “predicting” the composition of each annual sample as if it were a new observation.

To quantify accuracy and precision of our forecast predictions compared to observed genotypes, we used scale-dependent and scale-independent error metrics. For a scale-dependent metric we selected the widely used and easily understood mean squared error (MSE),

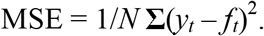

Where *N* = number of tests, *y_t_* = observed composition in the *t*^th^ test, and *f_t_* = the *t*^th^ forecast.

For a scale-independent metric we chose the less widely known mean arctangent absolute percentage error (MAAPE),

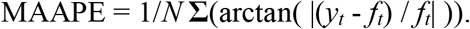

MAAPE can be interpreted intuitively as an absolute percentage error like the commonly used mean absolute percentage error (MAPE). However, MAAPE is less biased in greater penalties for positive errors than negative (Makridakis 1993). Moreover, the bounded range of the arctangent function (lim*_x_*_→∞_ tan^-1^ *x* = π/2) overcomes the MAPE’s limitation of going to infinity as the observed value approaches zero (Kim and Kim 2016).

For context, we compared the accuracy of our modeled estimates with interannual variability between members of paired samples (observed data) that had similar mean latitudes. We selected pairs of annual samples that differed by less than 0.25 degrees in mean latitude (2013 and 2014 at latitudes 43.8 and 43.5, respectively; and 2013 and 2015 at latitudes 43.8 and 43.9). This interannual variability should represent the minimum error possible in a forecast estimate. We used the R statistical package (R Core Team 2017) for most of our analyses and figures.

## RESULTS

NOAA fishery observers from the A-SHOP collected 9,862 individual tissue samples, which represented 41% of the total estimated Chinook Salmon bycatch between 2008 and 2015 (Table 1). Samples were stratified by year, and a total of 4,498 fish were randomly subsampled for DNA extraction and analysis (19% of total bycatch). Geographic distribution of individual tissue samples extended from Shelter Cove (41.43°), north to the Canadian border (48.48°); fishing depth 46 - 507 m (mean 246, SD 90.4); bottom depth 66 - 2,743 m (mean 427, SD 217.4), and fork length 24 - 113 cm (mean 58.7, SD 11.9). A slight female bias was observed (0.54). Mean latitude values for each annual bycatch sample ranged over 4.4 degrees (43.5**°** - 47.9**°)**. A general shift to the south was observed in fishing effort and bycatch beginning in 2011 (Table 1). Clearly, similar values for mean latitude might produce very different stock compositions if they have different distribution, e.g., non-normal. Through practical application and cross validation, our results demonstrate the sensitivity of our models to those violations of normality.

### Genotyping and genetic mixture modeling

Of 4,498 fish subsampled for genotyping, 96% met genotyping quality criteria. In addition to high genotyping success, high individual fish assignment probabilities were also observed. About 78% of fish successfully genotyped met the assignment probability criterion of ≥0.8 (N = 3,360, 14% of total bycatch; see Moran et al. 2014 for sensitivity analysis). Modeled genetic estimates of overall proportions derived from all fish successfully genotyped (“observed”) showed that most of the Chinook Salmon bycatch in this period came from Upper Klamath-Trinity River and S. Oregon and N. California Coastal ESUs (Fig. 1). Those two ESUs accounted for more than 50% of all bycatch in the study period. Secondary contributors included the Oregon Coast and Puget Sound ESUs and Southern British Columbia (lower Fraser River, data not shown).

**Figure 1.**
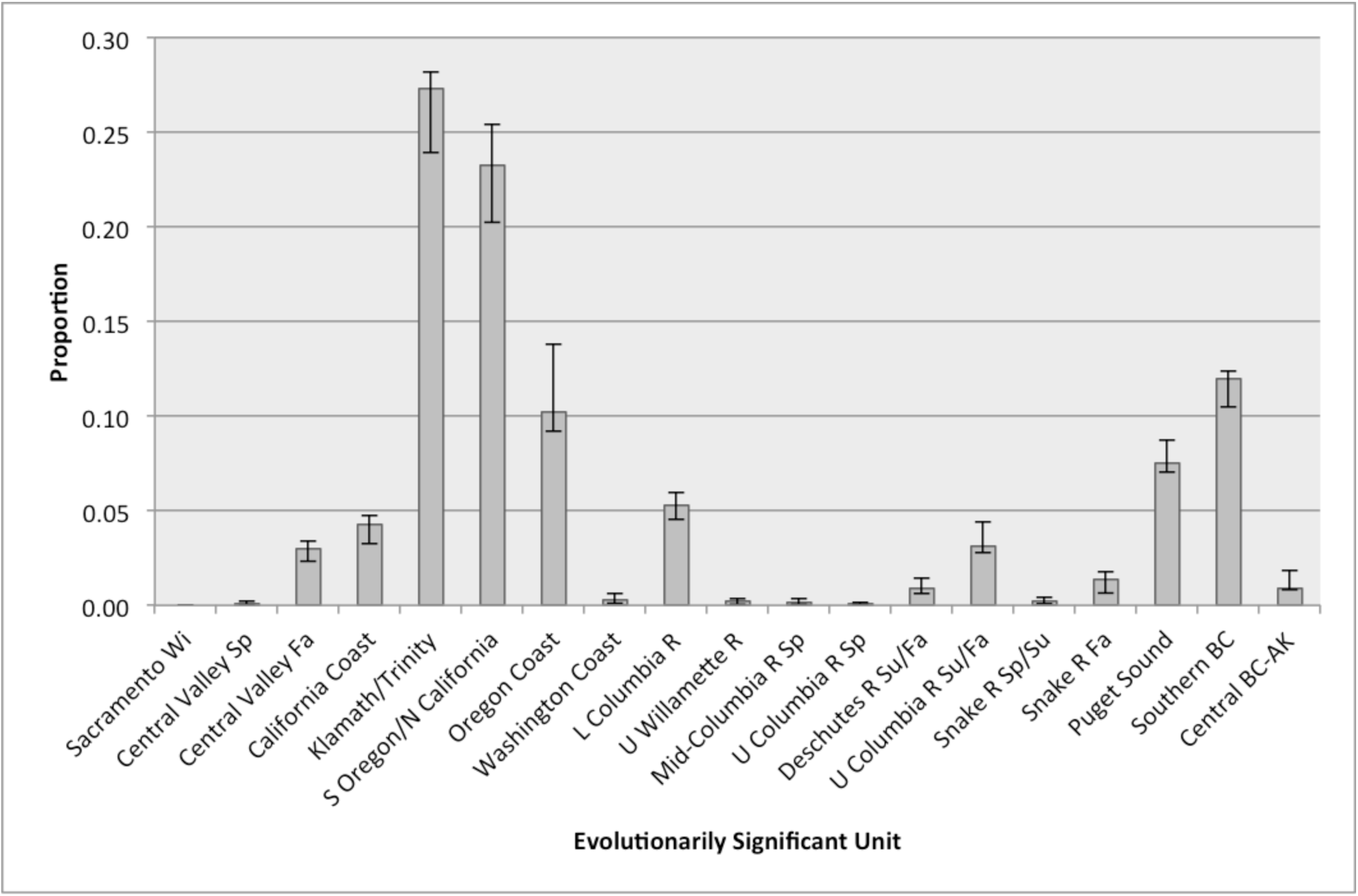
Overall Chinook Salmon ESU proportions observed in at-sea Pacific Hake trawl fisheries for all years combined 2008 - 2015. ESUs and two stock aggregates are ordered from south to north.

Managers often focus on fishery management areas. Differences in ESU composition between fishery management areas north and south of Cape Falcon (45.77**°)** showed the effect of latitude (See figs. 2 and 3 for area boundaries). The two southern ESUs—Upper Klamath-Trinity Rivers and S. Oregon/N. California—dominated bycatch south of Cape Falcon but dropped to only ∼5% each to the north of Cape Falcon. The opposite was true of the northern ESU, Puget Sound, and Southern BC populations. Spatial variation was also reflected when comparing the annual samples, which were each taken at different mean latitudes and showed different ESU proportions (see Jackknife cross validation below).

**Figure 2.**
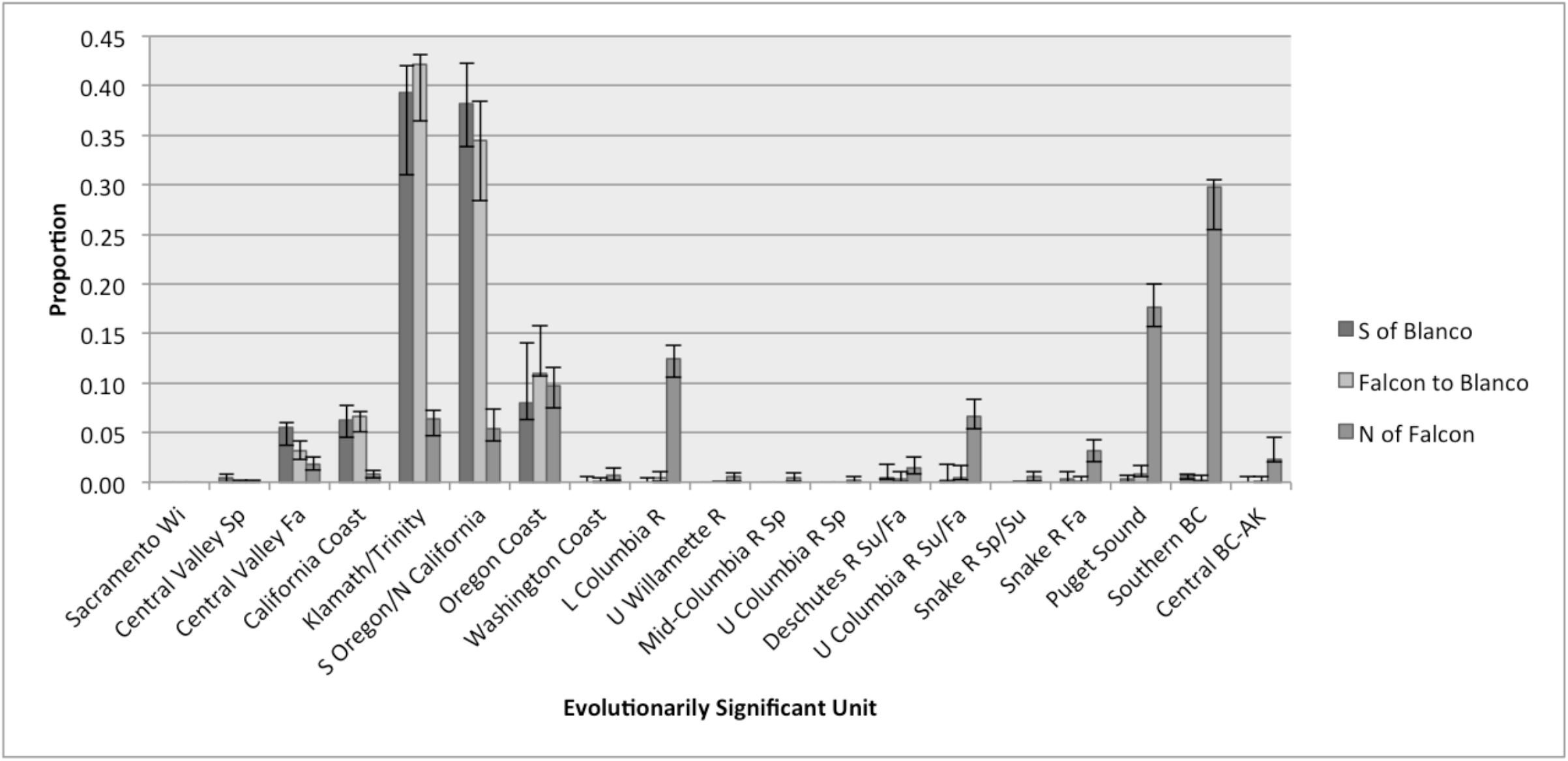
Chinook Salmon ESU proportions, stratified by fishery management area, ordered from south to north. Area 1: North of Cape Falcon, Oregon (45.77°); Area 2: Cape Falcon, Oregon to Cape Blanco, Oregon (42.75° to 45.77°); Area 3: Cape Blanco, Oregon to Cape Mendocino, California (40.16° to 42.75°). See Figure 3 for area boundaries.

**Figure 3.**
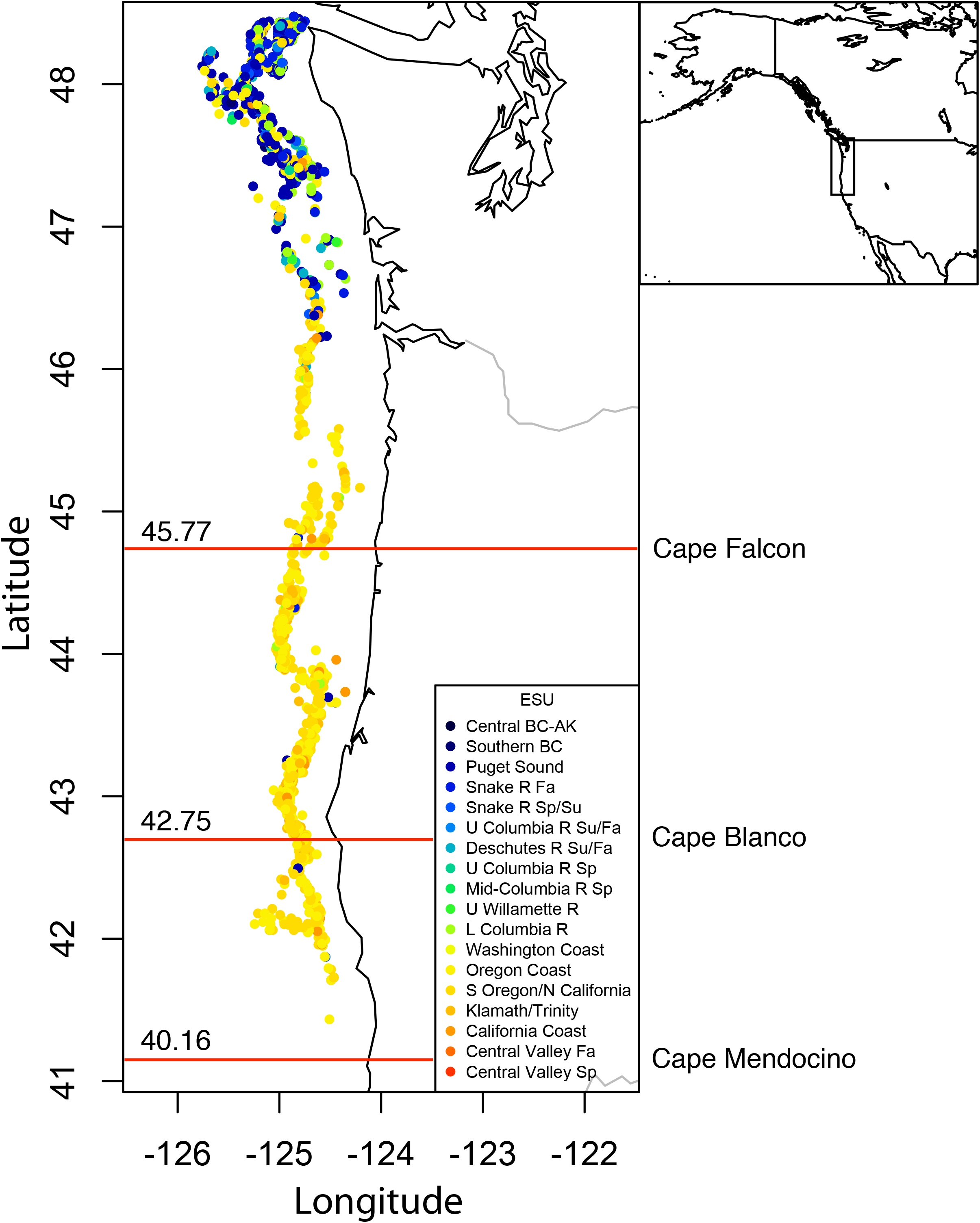
Individual Chinook Salmon taken in bycatch, color coded by most likely ESU of origin, from red in the south to blue in the north. Fishery management area boundaries are shown as red lines with associated latitudes (see Fig. 2 for area descriptions).

### Individual assignment

Again, our analysis included two fundamentally different approaches, modeled proportions versus individual assignment. The ESU stock composition results above describe fitted genetic models to all observed genotypic data simultaneously (Koljonen et al. 2005). At this point we shift to analyses based on individual fish assignment. First, we simply created a scatter plot of individually georeferenced fish, color coded by ESU, and overlaid on a map of the US West Coast (Fig. 3). That heuristic presentation also showed strong effects of latitude on ESU composition. No fish assigned to Sacramento Winter ESU, preventing inclusion of that ESU in the exploratory regression analysis described below.

Individual assignment was used to explore the ESU-specific predictive power of multiple sample attributes (e.g., latitude, depth, fork length, etc.). Previous simulations using the GAPS baseline showed that individual assignment with a threshold of ≥0.8 for inclusion, and regressing traits on ESUs is a robust and generally unbiased approach to inferring ESU-specific phenotypes (Moran et al. 2014). However, we recognize that individual assignment for compositional prediction raises issues of potential bias that go beyond the scope of this article. Nevertheless, our cross-validation approach subsumes those errors, allowing meaningful comparisons between disparate statistical methods. According to AIC evaluation of MLR models in particular, latitude was by far the most powerful single predictor of ESU origin (similar GAM results not shown). Other factors, most notably year, clearly explained additional variation, but Burnham and Anderson’s (2004) ΔAIC values for individual factors, relative to latitude alone, were compelling: year 1505, depth 1574, ordinal day 2271, and fork length 2544. More complex models gave lower AIC values, but were less suitable as predictors and as practical fishery management regulatory elements.

Plotting DR and MLR curves further illustrated the importance of latitude for most ESUs (Fig. 4). Consistent with the plot of fishery management areas (Fig. 2), northern ESUs were encountered primarily in the north, whereas southern ESUs were concentrated in southern and central coastal areas. Cape Falcon marked an abrupt transition in ESU composition of Chinook salmon bycatch.

**Figure 4.**
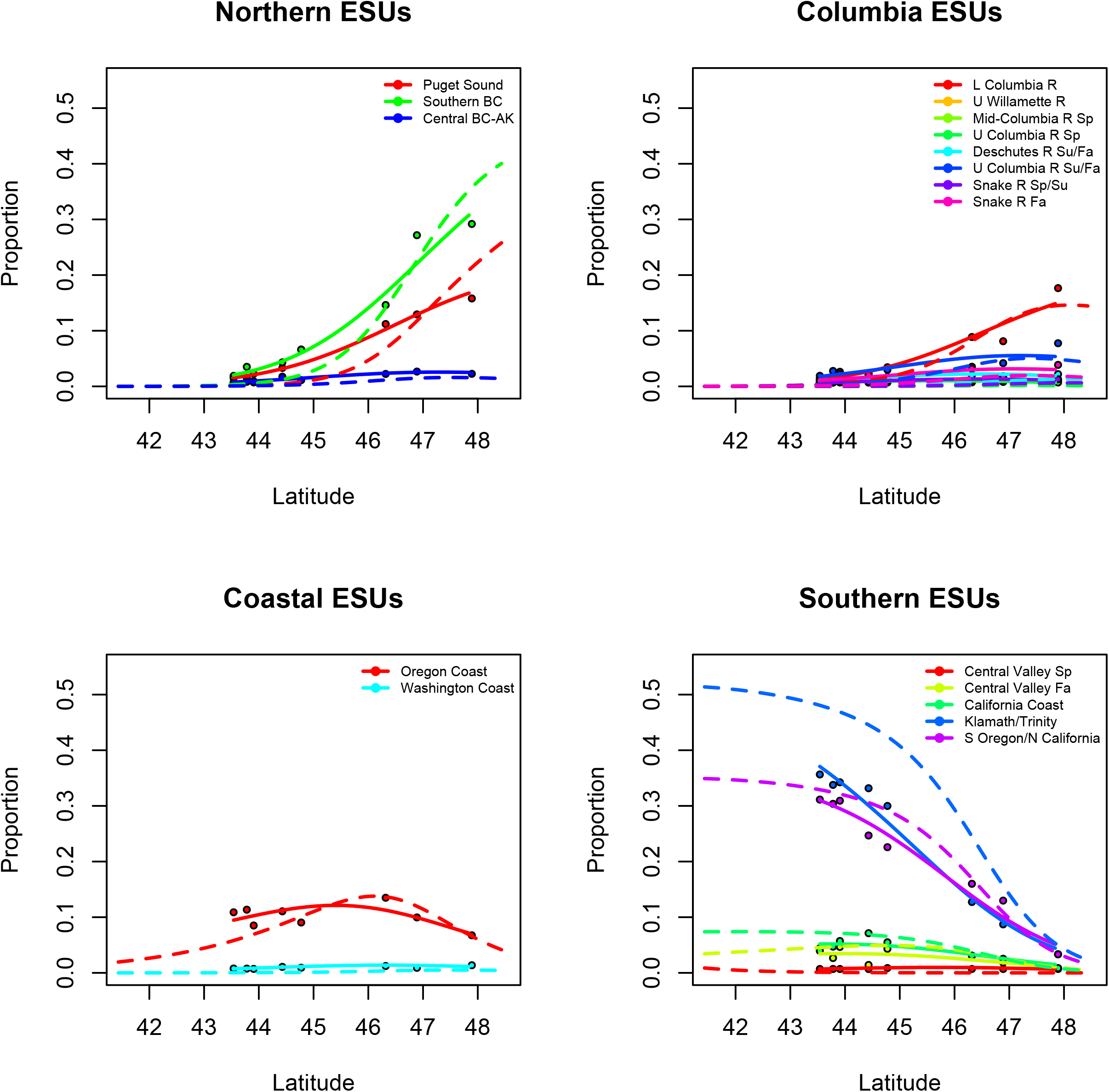
Dirichlet regression (solid) and multinomial logistic regression (dashed) with observed proportions (summing to one over all four panels) and mean latitudes of annual samples (points). Data ranges differ because DR is based on mean latitudes, whereas MLR is fitted to individual fish, their observed latitude, and the ESU to which they were assigned. Regional divisions simply facilitate interpretation.

### Jackknife cross validation

The estimated regression curves in Figure 4 can be compared with observed proportions in each annual sample (points in Fig. 4). However, the regression estimates include all data and are therefore not independent from the point observations in Figure 4 (i.e., potentially presenting an overly optimistic interpretation of predictive power). By contrast, the jackknife, cross-validation analysis provided an independent observation for every ESU proportion in every year. Figure 5 compares observed ESU proportions for each year with independently derived estimates from the two regression methods and latitude alone. Figure 6 almost perfectly summarizes all individual years (Fig. 4) by averaging over jackknife iterations. Despite a major difference in the MLR estimate for Upper Klamath/Trinity Rivers, estimates for both regression models were close to one another and close to observation for most ESUs.

**Figure 5.**
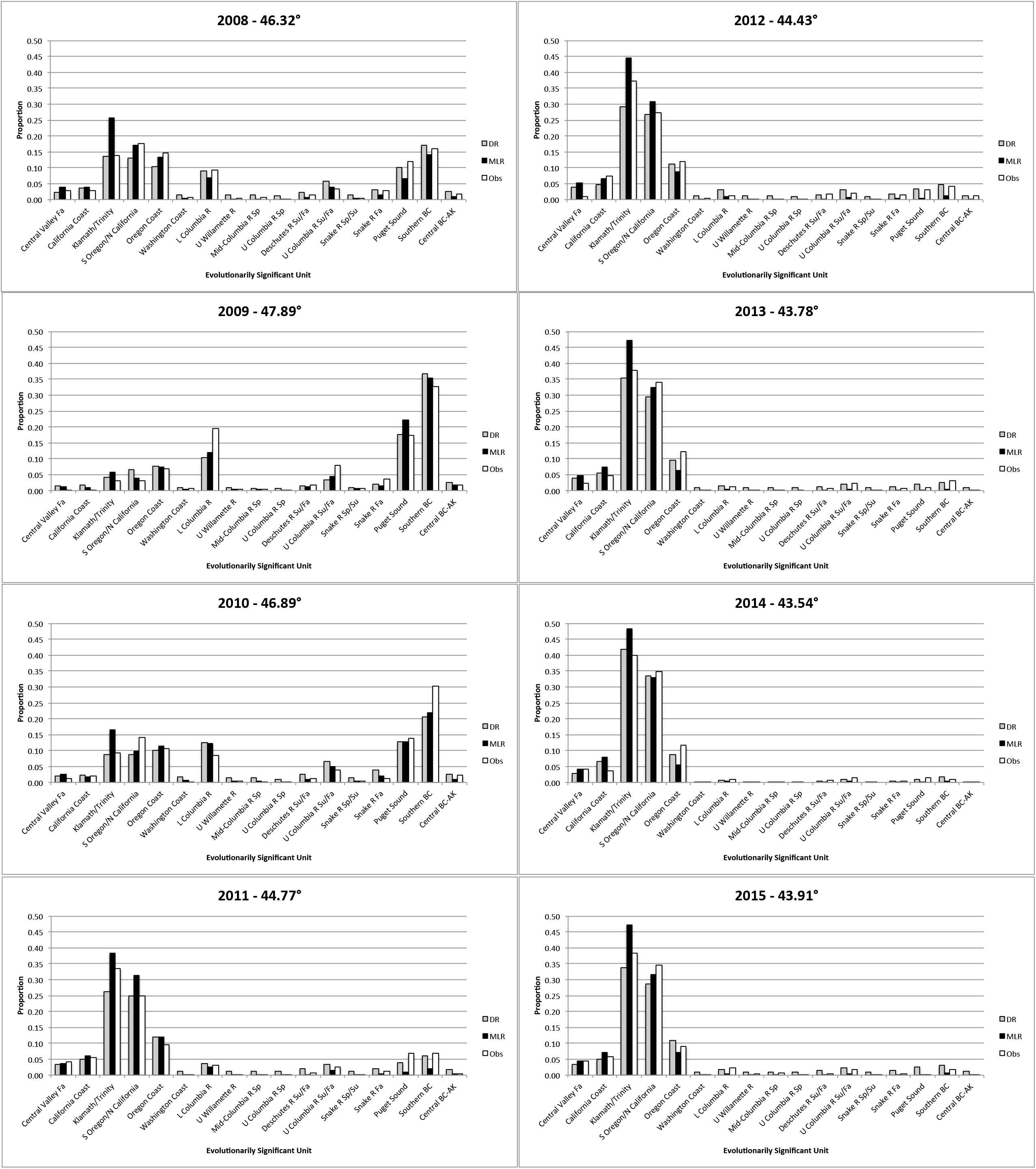
Selected realizations of the jackknife analysis are shown in each panel, where one entire annual sample (identified in title) was held out as a cross-validation test set (Obs). The remaining years were used as a training set to estimate ESU proportions given the observed latitude. Those proportions shifted beginning in 2011 and from 2012 to 2015 northern ESUs nearly disappeared. Note that observed proportions were similar between 2013 and 2015, when mean latitudes were also similar (∼43.8**°**, Table 1).

**Figure 6.**
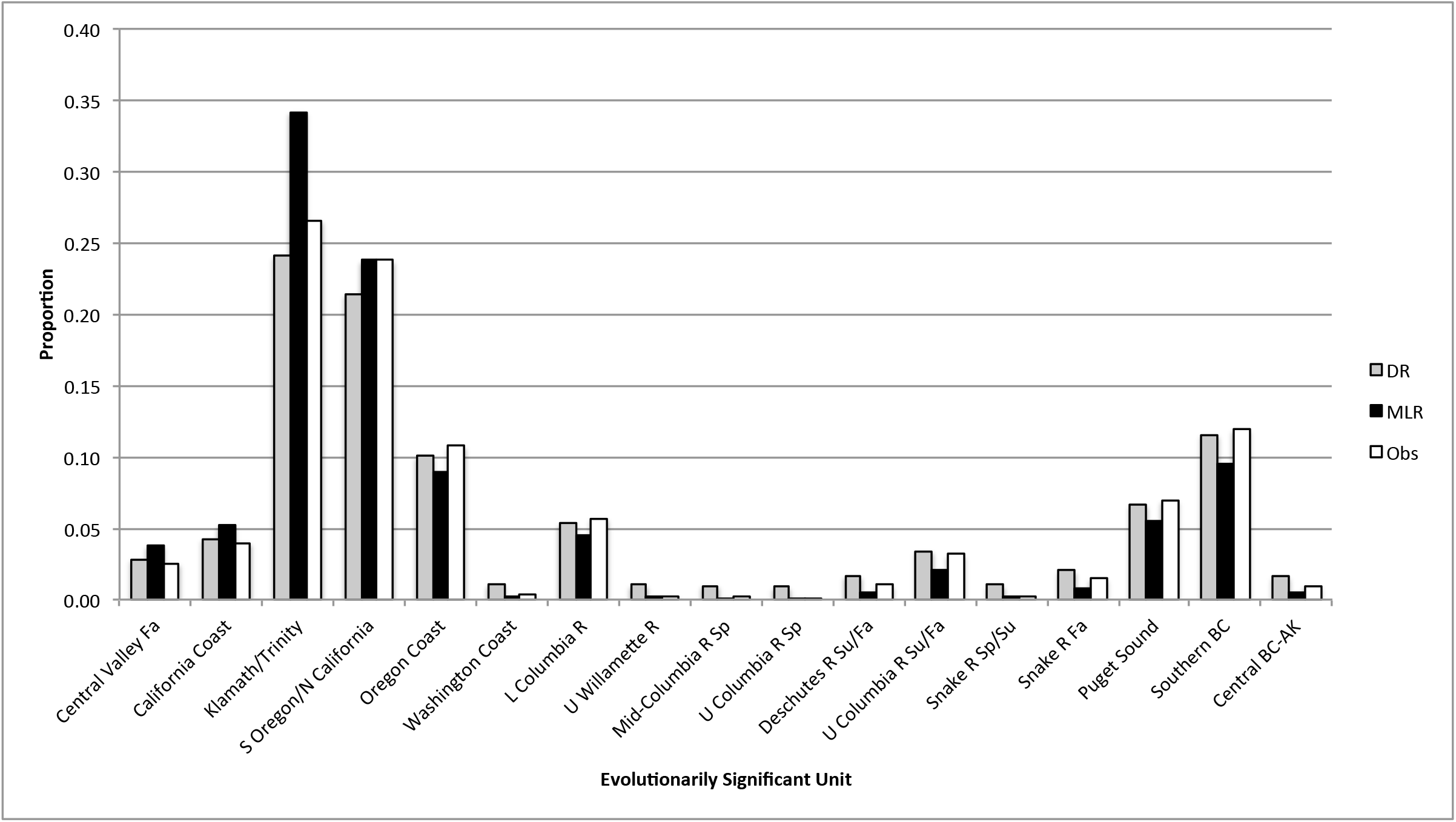
Model cross validation, summarized from Figure 5, comparing observed ESU proportions (Obs) with values predicted from Dirichlet regression (DR) and multinomial logistic regression (MLR), independent of observation.

### Annual replicates at similar latitude

By chance, some annual samples had very similar mean latitudes. Consistent with latitude being more important than year in the AIC, similar ESU proportions were observed in those annual samples that had similar mean latitudes (i.e., 2013, 2014, and 2015; Fig. 5). Comparison of variation between years with the accuracy of our forecast was mixed. Forecast predictions for large contributors showed substantially greater errors than variation observed between years at similar latitude (MSE = 0.0005 for DR and 0.0008 for MLR compared with only 0.0001 for the annual replicates). However, the scale-independent metric suggested that prediction errors were quite similar to interannual variation (MAAPE = 0.670 and 0.564 for DR and MLR versus 0.572 for interannual replicates).

### Accuracy of DR vs MLR, overall and by ESU

In predicting the ESU composition obtained from CMLMM in a given year, DR outperformed MLR for most ESUs that contributed more that 2.5%; however, the distribution of errors was complicated. For all contributors, MSE (most sensitive to large contributors) was 38% lower for DR than for MLR. Yet DR errors were more than three times as variable across years (CV = 56% for DR versus 21% for MLR). By contrast MAAPE (sensitive to small contributors) was 16% lower for MLR but nearly twice as variable (CV = 27% for MLR versus 14% for DR). So in each case, the more accurate regression method was less precise. A few patterns were evident in the accuracy and precision of individual ESU estimates. DR underpredicted Upper Klamath-Trinity Rivers ESU, in contrast to MLR that substantially overpredicted that ESU in nearly all years. Large MSE values for MLR were driven largely—but not entirely—by the extreme overprediction of Upper Klamath-Trinity Rivers ESU (Fig. 6; omitting Klamath/Trinity only reduced MSE from 0.0008 to 0.0007, still greater than 0.0005 for DR, and much greater than 0.0001 observed between years at the same latitude). With the notable exception of S. Oregon and N. California Coastal ESU, which MLR estimated almost exactly, southern ESUs were overestimated by MLR, whereas northern ESUs were underestimated. DR underestimated all larger contributors (>5%), but consistently overestimated smaller contributors, e.g., Washington Coast, Upper Willamette River, Mid-Columbia River Spring, Upper Columbia River Spring Snake River Spring/Summer, Snake River Fall ESUs [all years except 2009], and Central British Columbia and Alaska). Extensive additional analyses did not show an obvious effect of omitting fish that failed to meet the 0.8 probability criterion for MLR.

## Discussion

We succeeded in developing useful predictive models for Chinook Salmon ESU stock composition estimates in a fishery management context. Latitudinal clines and stock-specific distributions observed here were generally consistent with distribution of CWT recoveries in harvest fisheries (Weitkamp 2010; Shelton et al. 2019; Shelton et al. 2021). However, the current study, is the first we know of to describe pre-season prediction of ESU-specific impacts based simply on anticipated latitude of a proposed fishery. Given estimated bycatch numbers and latitudes (Matson and Erickson 2018), NOAA needed to parse total numbers by ESU to give predictions of actual fish counts, based on forecast proportions. These models were used previously in support of the 2017 Chinook salmon ITS and BiOp for the West Coast Groundfish Fishery Management Plan (NMFS—WCR 2017). This paper is an extension of the ground truthing that went into the BiOp. Because management needed to estimate ESU-specific impacts at various levels of take, our application required estimating proportions. Our approach was dictated by specific management needs, but the challenges of compositional forecasting and characterization of errors are ubiquitous in fields as disparate as ecological genetics and economic market research (Aitchison 1986). Even more broadly, the analysis of individuals versus aggregates is a fundamental statistical dichotomy.

### Broadly similar results between methods

We evaluated the performance of two accepted multinomial regression models representing two different analytical approaches, one based on genetic mixture modeling and DR, and the other based on individual assignment and MLR. We observed broadly similar results, despite differences in both the treatment of genetic data and in the regression. Independent cross validation showed that both analysis methods gave surprisingly accurate estimates despite a range of potential challenges. Not least, that different distributions of bycatch, with different stock compositions, might have similar means, thus confounding our prediction.

On initial inspection, Dirichlet regression appeared more accurate than MLR, but neither method provided a clear advantage across both rare and abundant ESUs. For example, MLR consistently overestimated the abundant Upper Klamath-Trinity Rivers ESU, especially in 2008 and 2011, whereas DR was much more accurate. DR was clearly more accurate than MLR in estimating large contributors; however, DR consistently overestimated small contributors, such as Upper Willamette River, Snake River Spring/Summer, and Snake River Fall—all listed as Threatened under the US Endangered Species Act. MLR was much more accurate than DR in estimating those small contributors at all latitudes and in nearly all years. Essentially all prediction errors for large contributors were greater than observed between members of paired annual samples at similar mean latitude—clearly room for improvement. However, for small contributors, prediction accuracy was extremely good. For example, 8 of 17 ESUs showed less variation between MLR prediction and observed ESU proportions in a given year than between years at similar latitude (6 of 17 for DR). In other words, prediction error with MLR was no larger than interannual variation for these small contributors.

Prediction accuracy is often critical for small contributors to mixed fisheries. Overprediction of sensitive or ESA-listed ESUs can lead to elevated concern, fishery restrictions, and needless forgone harvest. Moreover, a scale-independent metric like MAAPE could be considered a more appropriate measure of general forecasting accuracy than MSE. So, the recommended method depends on specific application and relative concern for abundant versus rare stocks. Harvest allocation might call for DR, whereas conservation might be better served by MLR. Managers are advised to examine both DR- and MLR-based take estimates and weigh the implications of specific inconsistencies.

### Comparison of Chinook Salmon stocks in bycatch versus directed harvest

We noted interesting differences from stock compositions previously reported for harvest. Direct comparison is difficult, but it appeared anecdotally that the Bellinger et al. (2015) study of commercial troll observed substantially higher proportions of Central Valley Fall ESU and lower proportions of Klamath-Trinity ESU than the current study. Also, Columbia River populations were more abundant in troll than in bycatch (Fig. 5 e, f, and g in Bellinger et al. 2015 compared to our Fig. 1). Similar results were seen in harvest studies (commercial and recreational fisheries) that sampled areas farther south (Winans et al. 2001; Satterthwaite et al. 2015).

With respect to latitude, the most direct available comparison between harvest and bycatch comes from the Washington commercial troll fishery (Moran et al. 2018). The mean latitude for all samples observed in that study between 2012 and 2015 was 47.4**°**, which was similar to the annual bycatch sample that we analyzed in 2009 (47.9**°**). Despite that similarity in latitude, the observed stock compositions were quite different. The relative abundance of Lower Columbia River and Upper Columbia River Summer/Fall was much higher in Washington troll than in bycatch from similar latitude. By contrast, proportions of Puget Sound and Southern British Columbia were much lower in troll than in bycatch. A difference worth noting is that at-sea bycatch tends to be unimodal in latitude within years (either north or south) but bimodal among years, whereas Washington coastal troll is strongly bimodal every year (Moran et al. 2018). There are also differences in depth and distance offshore. These anecdotal stock composition differences among disparate studies are difficult to interpret without broader spatial overlap and temporal replication. Eventually, however, the hope is that meta-analyses among fisheries can help discriminate where fish are caught from exactly where they are and how different populations use the marine environment in time and space.

### Fishery-dependent focus: Opportunities and limitations

Satterthwaite et al. (2014:128) pointed out that, for management purposes, “variability in interactions with the fishery is more relevant than ecological distribution.” The focus of the current study was not stock-specific, ecological distribution. Instead, we measured fishery impacts explicitly, and we limited most of our analysis to high-level spatial and temporal strata (i.e., coast-wide, annual). We chose both the scale and the compositional forecasting approach to provide fishery managers with a tool that required only simple inputs and would help meet obligations for predicting ESU impacts. In the context of depressed populations, it’s worth recognizing the inherent challenge in estimating very small bycatch numbers. Some ESUs were rare or absent in our analysis. Fewer than 10 fish were observed for each of six ESUs, and no Sacramento Winter fish were observed. Despite small numbers of observations for some ESUs, the expected proportions for those rare ESUs are not independent and are partially informed by the proportions of all other ESUs, including abundant ESUs for which estimates are much more confident. Despite concerns about interannual variability, non-normal latitudinal distributions, and the utility of mean latitude, our cross validation showed that error in the MLR estimate was essentially indistinguishable from interannual variation between samples taken at the same latitude. Again, we point out that “errors” in our regression estimates are relative to “observation” that is also an estimate but is based on observed genotypes and is our best estimates of ESU proportions in a given year. We use the term observation to emphasize the distinction between estimation and prediction, an observed mixed fishery versus a compositional forecast.

Results of Weitkamp (2010) suggested stable Chinook Salmon stock distributions across years, despite different ocean conditions and relative stock abundance (but see Satterthwaite et al. 2012; Shelton et al. 2019; Shelton et al. 2021). Seasonal variation in bycatch composition is substantially dampened, relative to directed harvest, because of the broad distribution of age classes in bycatch. For future genetic bycatch data, we will conduct a detailed exploration of stock-specific associations with oceanographic variables (sea-surface temperature, chlorophyll, etc.). Irrespective of spatial changes in fishing activity, if Chinook Salmon change their distribution or relative abundance in response to climate change, will the curves remain the same, shift uniformly north (or south), or will shapes and relative relationships of the curves change fundamentally, as predicted by Shelton et al (2021)? Climate effects on these latitudinal distributions could go beyond Chinook ESU impacts and conservation, to the extent ESA-listed southern resident killer whales target preferred Chinook salmon stocks (Hanson et al. 2010).

These questions are important, as the fishing industry becomes increasingly sophisticated in their requests for data on salmon distribution. Not only do fishing fleets want to avoid Chinook Salmon bycatch, they especially want to avoid protected or sensitive ESUs like Puget Sound, Lower Columbia River, Upper Klamath-Trinity Rivers, California Coast, and Central Valley Fall. Our figures 3 and 4 summarize what we know about where these ESUs are most likely to be caught. An obvious measure to avoid particular ESUs is to avoid nearby latitudes. The trouble is, moving north to avoid the threatened California Coast ESU, for example, would likely shift impacts to Puget Sound and Lower Columbia River, also listed as threatened under the US ESA. Moreover, moving fishing pressure north or south has a big effect on ESUs like Upper Klamath-Trinity Rivers or Puget Sound that have steep latitudinal clines, but less effect on ESUs like California Coast that have broader, flatter distributions. Despite that practical limitation to active avoidance of sensitive stocks, our regression models successfully address thorny statistical challenges and offer practical tools that are useful in evaluating ESU-specific impacts under different fishery management scenarios.

## ACKNOWLEDGEMENTS

We especially thank Andre Punt and William Satterthwaite of the PFMC’s Scientific and Statistical Committee. They questioned our early compositional analyses and started us down a very interesting and rewarding path. We thank Jeffrey Hard, Eric Ward, Jim Hastie, and Owen Hamel for helping develop the theoretical framework. NOAA’s A-SHOP and Cassandra Donovan provided outstanding training, sample collection, and data preparation. NWFSC laboratory staff Delia Patterson, Eric Iwamoto, Eric LaHood, and David Kuligowski conducted sample processing and genotyping. Many reviewers improved early drafts, including Michael Ford, Owen Hamel, Jeffrey Hard, Garrett McKinney, William Satterthwaite, Eric Ward, Laurie Weitkamp, and two anonymous reviewers. Reference to trade names is for information only, not endorsement by the U.S. Government.

## APPENDIX

Reference populations and reporting group structure for genetic mixture analysis based on Evolutionarily Significant Units (J. Myers, pers. comm. January 2016). Populations modified from Seeb et al. (2007). Status: E = Endangered, T = Threatened, C = Candidate, NW = Not Warranted, N/A = Not Applicable, stock aggregates that are not ESUs, which are only defined for the conterminous, US West Coast states. Carson Hatchery is a mixed-origin broodstock that is not listed under the ESA.

| Genetic baseline population | ESU reporting group | Status |
| --- | --- | --- |
| Sacramento Hatchery | Sacramento Winter | E |
| Butte Creek sp | Central Valley Spring | T |
| Deer Creek sp | Central Valley Spring | T |
| Mill Creek sp | Central Valley Spring | T |
| Feather Hatchery sp | Central Valley Fall | C |
| Feather Hatchery fa | Central Valley Fall | C |
| Butte Creek fa | Central Valley Fall | C |
| Stanislaus River | Central Valley Fall | C |
| Battle Creek | Central Valley Fall | C |
| Russian River | California Coastal | T |
| Eel River | California Coastal | T |
| Trinity Hatchery fa | Upper Klamath-Trinity Rivers | NW |
| Trinity Hatchery sp | Upper Klamath-Trinity Rivers | NW |
| Klamath River fa | Upper Klamath-Trinity Rivers | NW |
| Chetco River | S. Oregon and N. California Coastal | NW |
| Applegate Creek | S. Oregon and N. California Coastal | NW |
| Cole Rivers Hatchery | S. Oregon and N. California Coastal | NW |
| Elk Hatchery | Oregon Coast | NW |
| Sixes River | Oregon Coast | NW |
| Coquille River | Oregon Coast | NW |
| Coos Hatchery | Oregon Coast | NW |
| S Umpqua Hatchery | Oregon Coast | NW |
| Umpqua Hatchery | Oregon Coast | NW |
| Siuslaw River | Oregon Coast | NW |
| Alsea River | Oregon Coast | NW |
| Yaquina River | Oregon Coast | NW |
| Siletz River | Oregon Coast | NW |
| Salmon River fa | Oregon Coast | NW |
| Nestucca Hatchery | Oregon Coast | NW |
| Trask River | Oregon Coast | NW |
| Wilson River | Oregon Coast | NW |
| Kilchis River | Oregon Coast | NW |
| Nehalem River | Oregon Coast | NW |
| Necanicum Hatchery | Oregon Coast | NW |
| Forks Creek Hatchery | Washington Coast | NW |
| Humtulpis Hatchery | Washington Coast | NW |
| Queets River | Washington Coast | NW |
| Hoh River | Washington Coast | NW |
| Sol Duc Hatchery | Washington Coast | NW |
| Makah Hatchery | Washington Coast | NW |
| Lewis Hatchery sp | Lower Columbia River | T |
| Kalama Hatchery sp | Lower Columbia River | T |
| Cowlitz Hatchery sp | Lower Columbia River | T |
| Cowlitz Hatchery fa | Lower Columbia River | T |
| Sandy River | Lower Columbia River | T |
| Lewis River fa | Lower Columbia River | T |
| Spring Creek Hatchery | Lower Columbia River | T |
| McKenzie Hatchery | Upper Willamette River | T |
| N Santiam Hatchery | Upper Willamette River | T |
| Warm Springs Hatchery | Mid-Columbia River Spring | NW |
| John Day River | Mid-Columbia River Spring | NW |
| U Yakima Hatchery | Mid-Columbia River Spring | NW |
| Wenatchee River sp | Upper Columbia River Spring | E |
| Wenatchee Hatchery sp | Upper Columbia River Spring | E |
| Carson Hatchery | Upper Columbia River Spring | N/A |
| U Deschutes River | Deschutes River Summer/Fall | NW |
| L Deschutes River | Deschutes River Summer/Fall | NW |
| Hanford Reach | Upper Columbia River Summer/Fall | NW |
| Wenatchee River su/fa | Upper Columbia River Summer/Fall | NW |
| Wells Hatchery | Upper Columbia River Summer/Fall | NW |
| Methow River | Upper Columbia River Summer/Fall | NW |
| Lyons Ferry Hatchery | Snake River Fall | T |
| EF Salmon River | Snake River Spring/Summer | T |
| WF Yankee Fork | Snake River Spring/Summer | T |
| Secesh River | Snake River Spring/Summer | T |
| Rapid River Hatchery | Snake River Spring/Summer | T |
| Minam River | Snake River Spring/Summer | T |
| Imnaha River | Snake River Spring/Summer | T |
| Newsome Creek | Snake River Spring/Summer | T |
| Tucannon Hatchery | Snake River Spring/Summer | T |
| Tucannon River | Snake River Spring/Summer | T |
| Clear Creek Hatchery | Puget Sound | T |
| Voights Hatchery | Puget Sound | T |
| S Prairie Creek | Puget Sound | T |
| Soos Hatchery | Puget Sound | T |
| George Adams Hatchery | Puget Sound | T |
| Hamma Hamma River | Puget Sound | T |
| Snoqualmie River | Puget Sound | T |
| Samish Hatchery | Puget Sound | T |
| Elwha Hatchery | Puget Sound | T |
| Elwha River | Puget Sound | T |
| Dungeness River | Puget Sound | T |
| NF Nooksack Hatchery | Puget Sound | T |
| White Hatchery | Puget Sound | T |
| Hatcheryupp Sp Hatchery | Puget Sound | T |
| Skykomish River | Puget Sound | T |
| Wallace Hatchery | Puget Sound | T |
| NF Stillaguam Hatchery | Puget Sound | T |
| Skagit River | Puget Sound | T |
| U Sauk River | Puget Sound | T |
| Suiattle River | Puget Sound | T |
| L Sauk River | Puget Sound | T |
| Marblemount Hatchery sp | Puget Sound | T |
| Marblemount Hatchery su | Puget Sound | T |
| U Cascade River | Puget Sound | T |
| U Skagit River | Puget Sound | T |
| W Chilliwack Hatchery | Southern BC | N/A |
| Maria Slough | Southern BC | N/A |
| Birkenhead Hatchery | Southern BC | N/A |
| M Shuswap Hatchery | Southern BC | N/A |
| L Thomson River | Southern BC | N/A |
| L Adams Hatchery | Southern BC | N/A |
| Clearwater River | Southern BC | N/A |
| Riveraft River | Southern BC | N/A |
| Spus Hatchery | Southern BC | N/A |
| Nicola Hatchery | Southern BC | N/A |
| Louis Creek | Southern BC | N/A |
| Deadman Hatchery | Southern BC | N/A |
| U Chilcotin River | Southern BC | N/A |
| Chilko River | Southern BC | N/A |
| Quesnel River | Southern BC | N/A |
| Nechako River | Southern BC | N/A |
| Stuart River | Southern BC | N/A |
| Swift River | Southern BC | N/A |
| Morkill River | Southern BC | N/A |
| Salmon River sp | Southern BC | N/A |
| Cowichan Hatchery | Southern BC | N/A |
| Nanaimo Hatchery fa | Southern BC | N/A |
| Big Qualicum Hatchery | Southern BC | N/A |
| Puntledge Hatchery fa | Southern BC | N/A |
| Quinsam Hatchery | Southern BC | N/A |
| Nitinat Hatchery | Southern BC | N/A |
| Sarita Hatchery | Southern BC | N/A |
| Tranquil River | Southern BC | N/A |
| Robertson Hatchery | Southern BC | N/A |
| Conuma Hatchery | Southern BC | N/A |
| Tahsis River | Southern BC | N/A |
| Marble Hatchery | Southern BC | N/A |
| Porteau Cove Hatchery | Southern BC | N/A |
| Klinaklini River | Southern BC | N/A |
| Wannock Hatchery | Central BC-AK | N/A |
| Atnarko Hatchery | Central BC-AK | N/A |
| Kitimat Hatchery | Central BC-AK | N/A |
| Ecstall River | Central BC-AK | N/A |
| L Kalum River | Central BC-AK | N/A |
| Bulkley River | Central BC-AK | N/A |
| Sustut River | Central BC-AK | N/A |
| Kincolith River | Central BC-AK | N/A |
| Kwinageese River | Central BC-AK | N/A |
| Damdochax River | Central BC-AK | N/A |
| Owegee River | Central BC-AK | N/A |
| L Tahltan River | Central BC-AK | N/A |
| Nakina River | Central BC-AK | N/A |
| Kowatua Creek | Central BC-AK | N/A |
| Tatsatua Creek | Central BC-AK | N/A |
| U Nahlin River | Central BC-AK | N/A |
| Chickamin/White Hatchery | Central BC-AK | N/A |
| Chickamin Hatchery | Central BC-AK | N/A |
| Chickamin River | Central BC-AK | N/A |
| Keta River | Central BC-AK | N/A |
| King Creek | Central BC-AK | N/A |
| Clear Creek | Central BC-AK | N/A |
| Creekipple Creek | Central BC-AK | N/A |
| Andrew Creek | Central BC-AK | N/A |
| Andrew/Mac Hatchery | Central BC-AK | N/A |
| Andrew/Med Hatchery | Central BC-AK | N/A |
| Andrew/Cry Hatchery | Central BC-AK | N/A |
| King Salmon River | Central BC-AK | N/A |
| Big Boulder Creek | Central BC-AK | N/A |
| Tahini River | Central BC-AK | N/A |
| Tahini/Mac Hatchery | Central BC-AK | N/A |
| Klukshu River | Central BC-AK | N/A |
| Situk River | Central BC-AK | N/A |

<sup>1</sup>Feather River spring run brood stock was extensively hybridized with fall run in that program and the two are now genetically indistinguishable.
<sup>2</sup>Brood stock from Cole M. Rivers Hatchery on the Rogue River in Southern Oregon is currently propagated and released in Young's Bay at the mouth of the Columbia River.

## Footnotes

1 Also referred to by the industry name “Pacific whiting”

